# A streamlined workflow for automated cryo focused ion beam milling

**DOI:** 10.1101/2020.02.24.963033

**Authors:** Sebastian Tacke, Philipp Erdmann, Zhexin Wang, Sven Klumpe, Michael Grange, Jürgen M. Plitzko, Stefan Raunser

## Abstract

Cryo-electron tomography is an emerging technique to study the cellular architecture and the structure of proteins at high resolution *in situ*. Most biological specimens are too thick to be directly investigated and are therefore thinned by milling with a focused ion beam under cryogenic conditions. This procedure is prone to frost and amorphous ice depositions which makes it a tedious process, leading to suboptimal results especially when larger batches are milled. Here, we present new hardware that overcomes the current limitations. We developed a new glove box and a high vacuum cryo transfer system and installed a stage heater, a cryo-shield and a cryo-shutter in the FIB milling microscope. This tremendously reduces the ice depositions during transfer and milling, and simplifies the handling of the sample. In addition, we tested a new software application that automates the key milling steps. Together, these improvements allow for high-quality, high-throughput cryo-FIB milling.

## Introduction

Electron tomography at cryogenic temperatures (cryo-ET) offers the unique possibility to structurally analyse biological macromolecules in their native cellular environment [Grange2017, Bartesaghi2012]. Using this method, it is even possible to determine protein structures at near-atomic resolution by applying sub-volume averaging techniques [Schur2016].

One prerequisite for electron cryo microscopy (cryo-EM) is that the sample is thinner than the inelastic mean-free-path length of electrons (approximately 350 nm for 300 keV electrons [Vulovic2013]. Thicker samples result in an increased number of inelastically scattered electrons, which decreases the quality of the image. Unfortunately, the majority of biological samples (e.g. eukaryotic cells or tissues) are too thick for cryo-ET and must therefore be thinned.

Cryo-ultramicrotomy is one approach that can be used to reduce specimen thickness. Here, the sample is mechanically cut into ribbons of vitreous sections that are thin enough to be imaged by cryo-ET [Al-Amoudi2004]. However, this method is technically demanding and prone to sample-altering artefacts such as compression, knife marks or crevasses [Al-Amoudi2005]. Another approach uses a focused ion beam under cryogenic conditions (cryo-FIB) in a scanning electron microscope (SEM) to thin samples [Marko2007, Rigort2010]. In recent years, numerous studies have demonstrated the suitability of this technique for preparing biological samples for cryo-ET without the artefacts known from cryo-ultramicrotomy [Hagen2015, Mahamid2016].

Current best-practice protocols for cryo-FIB milling include: i) vitrification of the sample, ii) identification of regions-of-interest (ROI), iii) rough ablation of the surrounding sample material and iv) polishing of the target region, leaving a thin lamella ready for cryo-ET [Schaffer2017]. However, the entire procedure involves multiple handling, transfer, milling and imaging steps. During these steps, samples need to be handled with forceps and it is difficult even for highly experienced practitioners not to destroy the thin and fragile lamellae. Moreover, since the vitrified sample needs to be cooled constantly to avoid a phase transition to hexagonal ice (devitrification temperature around 135 K), it is especially prone to crystalline and amorphous ice contamination. Contamination of polished lamellae is particularly disruptive to cryo-ET as ice crystals often obscure the view on features otherwise visible in the lamellae. It is even possible that ice crystals (frost contamination) conceal the entire lamella, which prevents any meaningful data acquisition (Figure 1a). Besides the transfer, which bears a high risk of exposing the sample to water, a potential source of (amorphous) ice contamination is the cryo-FIB/SEM itself. Despite the high vacuum inside the SEM, water molecules from the residual gas inside the microscope chamber can still deposit on the sample that is kept well below −140 °C to avoid devitrification. Depending on the partial pressure of water inside the chamber, this effect can be severe and becomes especially problematic when samples reside in the SEM for several hours (Figure 1b,c).

**Figure 1:**
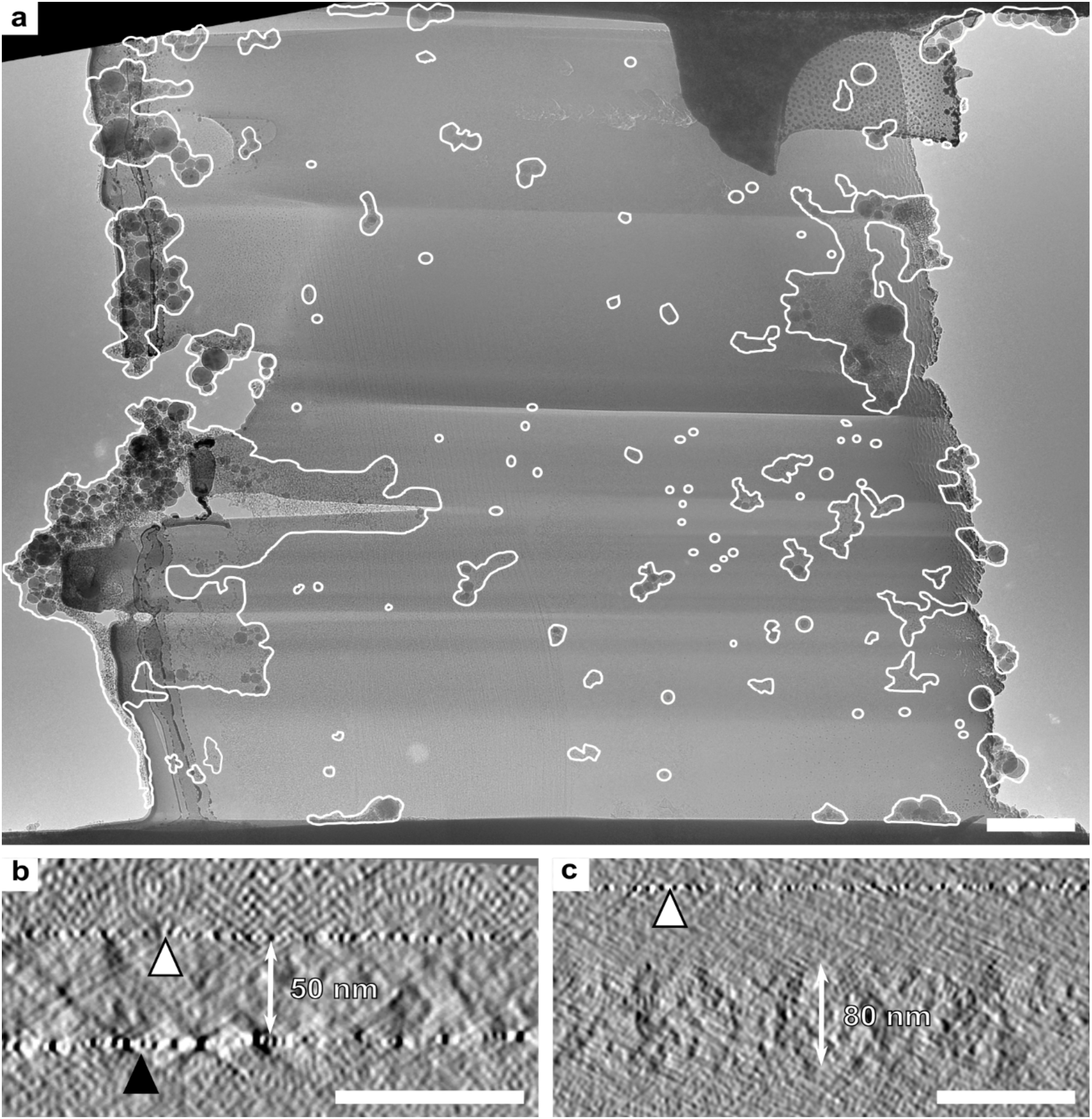
Possible artefacts arising during handling, transfer and processing. **a**) Overview of a lamella, which was prepared under standard conditions without hardware upgrades. When the sample transferred in a hydrous environment, ice crystals of different sizes (highlighted by marked areas) contaminate up to 20% of the lamella area. Scale bar, 1 μm. **b,c**) Side views of two tomograms, taken from two different lamellae, prepared in the same milling session. The lamella in (**b**) was polished last. Thus, it was taken out of the SEM almost directly after polishing. Consequently, the water deposition is not detectable. The sputter layer (white triangle) is directly on top of the lamella. The black triangle marks some platinum re-deposition. The lamella in (**c**) was polished approximately 1 hour before the lamella in (**b**) and stayed the whole time in the SEM. In this case, a ~50 nm thick water deposition can be identified on top of the 80 nm thick lamella, reducing its quality. The platinum coating above the sample material is indicated by a white triangle. Scale bars, 100 nm.

Recently, several software packages that automate the milling process were introduced by ThermoFisher, Zeiss [Zach2020] and the DeMarco group [Buckley2020]. The automation enables the batch milling of up to 20-27 lamellae in 24 h and reduces the time the operator has to spent actively at the SEM. However, during this process the samples remain for many hours in the microscope, which considerably increases contamination and limits the number of lamellae with sufficiently good quality per session.

To obtain optimal lamellae for cryo-ET and to unfold the full strength of automated cryo-FIB milling, the following major hurdles need to be overcome: First, all types of frost contamination need to be reduced to a minimum. Second, all amorphous ice contamination inside the microscope needs to be avoided. Third, the handling of the lamellae must be streamlined to avoid lamella cracking.

Here, we present an integrated workflow that addresses and overcomes all three bottlenecks. In a first step, we implemented a specially designed glove box and a high vacuum cryo transfer system into the current workflow to reduce frost contamination during handling and transfer. Secondly, we tremendously reduced the contamination rates inside the cryo-FIB/SEM by the installation of a stage heater, a cryo-shield and a cryo-shutter. Thirdly, we implemented new tools to simplify the sample preparation and handling.

In addition, we tested the new commercial software application AutoTEM Cryo (ThermoFisher). We show that in combination with this software, the new hardware increases the overall throughput of high-quality lamella production, enabling new types of experiments, which are considered infeasible with current setups.

## Results

### Sample preparation and handling

After vitrification by plunge or high-pressure freezing, the EM grid with the sample is placed into the Autogrid, then put into the Autogrid shuttle and transferred to the cryo-FIB/SEM. During these steps, the sample needs to be kept at temperatures below −140°C to avoid devitrification. Consequently, the sample is prone to any kind of contamination. To reduce frost contamination, many cryo-EM laboratories have implemented air-dehumidifier rooms. While those help to reduce the overall humidity, and thereby frost contamination, they cannot completely prevent it.

To reduce frost contamination during handling and simplify implementation of the technique in laboratories unable to control room humidity, we developed a glove box system (Figure 2a) that allows the grid preparation steps to be performed in a tightly regulated atmosphere. During operation, the glove box is constantly purged with dry nitrogen gas (10 ppm H_2_0) to keep the humidity below the detection limit (1 %) of the humidity sensors used. Beyond functionality, the glove box system was designed to provide maximal convenience during grid preparation. To guarantee that the sample can be brought into the box without disturbing the anhydrous environment inside, we equipped the glove box with a load lock and a port for a high vacuum cryo transfer system. The direct supply of liquid nitrogen is provided by a liquid nitrogen reservoir, which is located inside the glove box. Furthermore, we installed a set of heated tool holders (Figure 2a, marked area) in the box that allow to heat and dry different tools, such as forceps, clipping tools, cassette gripper, etc. directly after usage.

**Figure 2:**
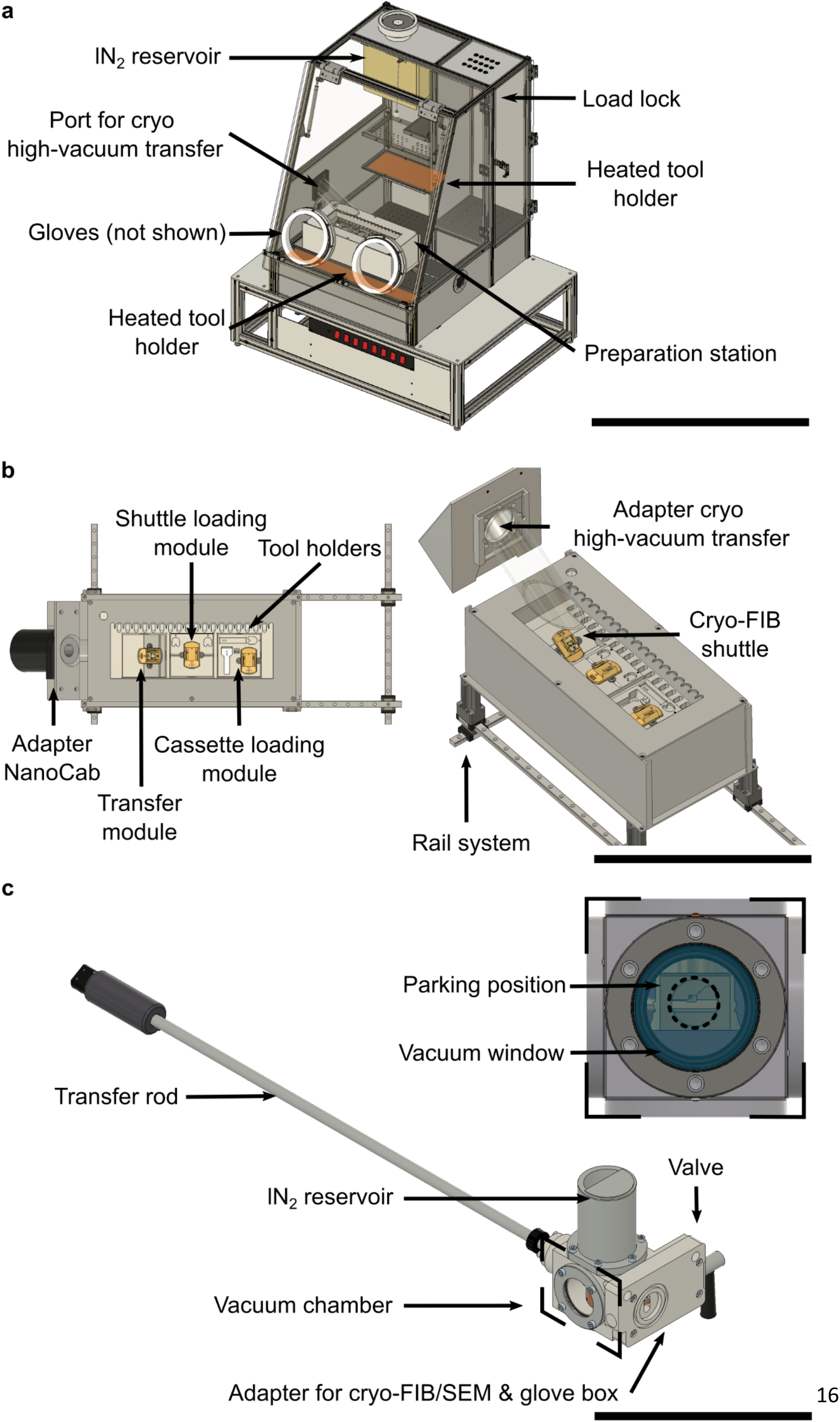
Schematic of the glove box, the preparation station, and the high vacuum cryo transfer system. **a**) The glove box enables convenient sample handling in an anhydrous environment. To reduce the humidity to a minimum, the glove box is purged during operation with nitrogen and a cold trap is installed in the back of the lN_2_ reservoir. lN_2_ can be taken from this reservoir for applications inside the glove box. For convenient handling, heated tool holders are installed. A load lock enables the insertion of tools without breaking the humidity barrier. Scale bar, 600 mm. **b**) A new preparation station guarantees a comfortable and secure handling of the sample material. The preparation station consists of four parts: the liquid nitrogen bath, the preparation modules, the adapter for the transfer systems and the adapter for the NanoCab. All modules can be re-arranged inside the liquid nitrogen bath according to the preferences of the operator and are designed for left- and right-handed persons. Clipped autoloader C-clip rings can be loaded onto the cryo-FIB shuttle or to the autoloader cassette using the corresponding modules. Also clipping is possible in the preparation station but is not shown in this overview. The preparation station is mounted onto a rail system to ensure highest flexibility inside the glove box. Scale bar, 400 mm. **c**) The high vacuum cryo transfer system consists of a transfer rod for picking up the cryo-FIB shuttle, a vacuum chamber, liquid nitrogen reservoir, a valve and the adapter for the cryo-FIB/SEM and glove box. The sample can be picked up by the transfer rod and is inserted in the vacuum chamber. Inside the vacuum chamber the cryo-FIB shuttle is placed in a parking position. Here, the shuttle is actively cooled by the lN_2_ reservoir. The liquid nitrogen acts also as a cryo-pump and maintains the vacuum level, while the high vacuum cryo transfer is disconnected from the pumps. The lN_2_ volume holds for approximately 30 min. After the shuttle is positioned in the parking position, the valve towards the glove box is closed and the high vacuum cryo transfer is evacuated. Thereafter, the valve at the transfer system is closed and the sample can be transferred to the cryo-FIB/SEM. Scale bar, 300 mm.

To increase the ease of handling grids inside the box, we designed and built a dedicated universal preparation station (Figure 2b), again maximising the convenience for the user. The station contains an isolated liquid nitrogen bath with three different preparation modules for each step of grid preparation, namely autoloader grid assembly, cryo-FIB shuttle loading and autoloader cassette loading. The users can arrange the needed modules according to their working routine.

To avoid amorphous ice contamination and devitrification during sample transfer from the glove box to the cryo-FIB/SEM, we implemented a previously developed high vacuum cryo transfer system [Tacke2016] (Figure 2c), which consists of a transfer rod, a liquid nitrogen reservoir, a vacuum chamber, a valve and a plug & play adapter for the glove box or cryo-FIB/SEM. To avoid devitrification during transfer, the cryo-FIB shuttle is actively cooled by the liquid nitrogen reservoir as soon as the transfer rod is retracted in its parking position. Moreover, because the cooled surfaces of the nitrogen reservoir act as a cryo-pump, the vacuum level is kept constant as long as the reservoir is filled [Tacke2016].

With the introduced devices, we are able to minimise the adverse implications, currently associated with existing workflows streamlining the entire grid preparation whilst improving user convenience, efficiency and flexibility. Especially during sample screening, where many samples have to be prepared and transferred over a longer timeframe, the benefit of the devices is apparent. Due to the reduced humidity, the preparation station can stay cooled for several hours without noticeable frost contamination and the high vacuum cryo transfer system allows the necessary quick and contamination-free transfer (Supplementary Video).

### Hardware modification of the cryo-FIB

Although cryo-FIB/SEMs are typically operated at high vacuum conditions, contamination of the sample inside the microscope cannot be completely avoided, due to various sources of contamination in the FIB chamber. One of these is the ablated sample material which re-deposits on the specimen. This kind of contamination is greatest at the beginning of lamellae preparation when the sample material is milled in bulk and decreases to a minimum towards the final polishing step. It is therefore not a major problem for the production of lamellae if the milling is performed in two stages: first all sites are roughly milled to a thickness of approximately 500 - 800 nm and only then the final polishing is performed.

The second source of contamination is residual water inside the vacuum chamber. Its deposition/sublimation rate depends on two factors: i) the fraction of water in the residual gas mixture (partial pressure) and ii) the surface temperature of the sample material (Figure 3a). We installed a large cryo-shield and a cryo-shutter to reduce the partial pressure of water in the system to less than 4 x 10^−9^ mbar and heated the sample to max. −165 °C using a newly installed stage heater (Figure 4a, Figure 4b). Combined, these actions moved the ratio between deposition and sublimation closer towards an equilibrium, i.e. minimal deposition rates, which is optimal for cryo-FIB milling (Figure 3a) (see also [Umrath1983], [Echlin1992]).

**Figure 3:**
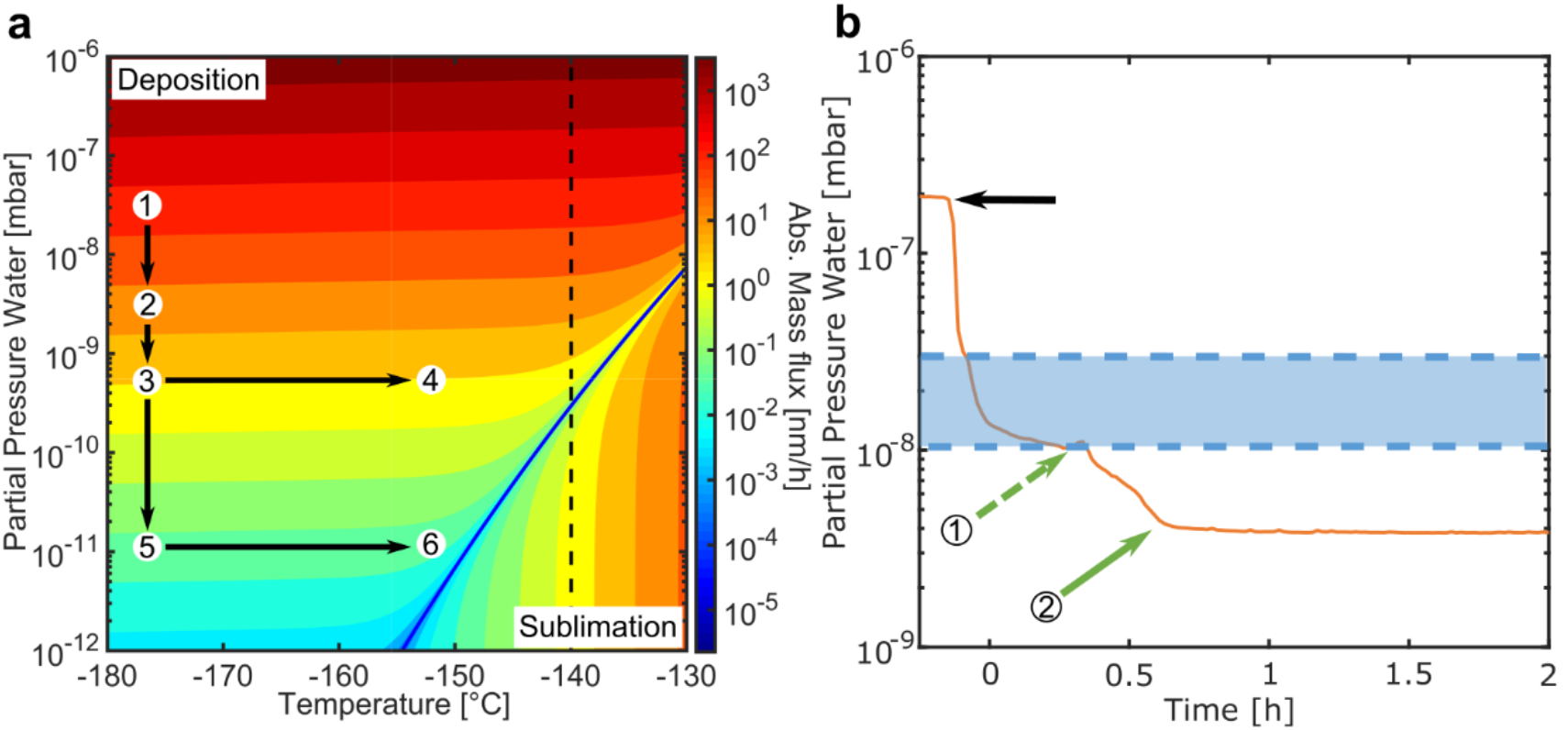
Phase diagram of water. **a**) Phase diagram of water and the associated mass flux (deposition/sublimation) [Umrath1983]. The encircled numbers mark the different measurement conditions. The dashed line indicates the devitrification temperature of water. The arrows indicate our strategy to minimize the ice deposition (amorphous ice contamination) onto the sample. After the installation of the cryo-shield, cryo-shutter Type I and stage heater, the contamination rate increases from the specified 50 nm h^−1^ to 85 ± 7 nm h^−1^ (condition 1). As soon as the cryo-shield is cooled, the partial pressure of water drops to 4 x 10^−9^ mbar and the contamination rate decreases to 72 ± 4 nm h^−1^ (condition 2). When the cryo-shutter Type I is inserted, the partial pressure in the vicinity of the sample decreases by approximately one order of magnitude (condition 3), which reduces the contamination rate to 12.8 ± 1.6 nm h^−1^. To reduce the contamination rate even further, the sample is heated to approximately −165 °C. Since there might be a temperature difference of 15 °C between the stage and the shuttle, the temperatures for condition 4 and 5 are specified as approximately −150 °C. The heating reduces the contamination rate to 5.6 ± 1.0 nm h^−1^ (condition 4), which is already a tremendous improvement compared to the initial contamination rate of 50 nm h^−1^. To reduce the contamination even further we designed the more effective cryo-shutter Type II (Supplementary Figure 1). Using this shutter, we were able to reduce the contamination to a non-detectable level (−0.6 ± 1.3 nm h^−1^) (condition 6), even without heating the sample (0 ± 2 nm h^−1^) (condition 5). **b**) Measured partial pressure of water inside the cryo-FIB/SEM chamber (orange graph). As soon as the SEM and cryo-shield are cooled (black arrow), water molecules from the residual gas get trapped and the partial pressure decreases. After 15 min, the stage reaches its end temperature of −185°C (dashed green arrow). Under this condition, the impact of the cryo-shield is still neglectable, since its temperature is still above −100°C. After 30 min, also the cryo-shield reaches its end temperature and the system reaches its final partial pressure value (solid green arrow). The blue area indicates the partial pressure regime of standard cryo-FIB/SEM instruments.

**Figure 4:**
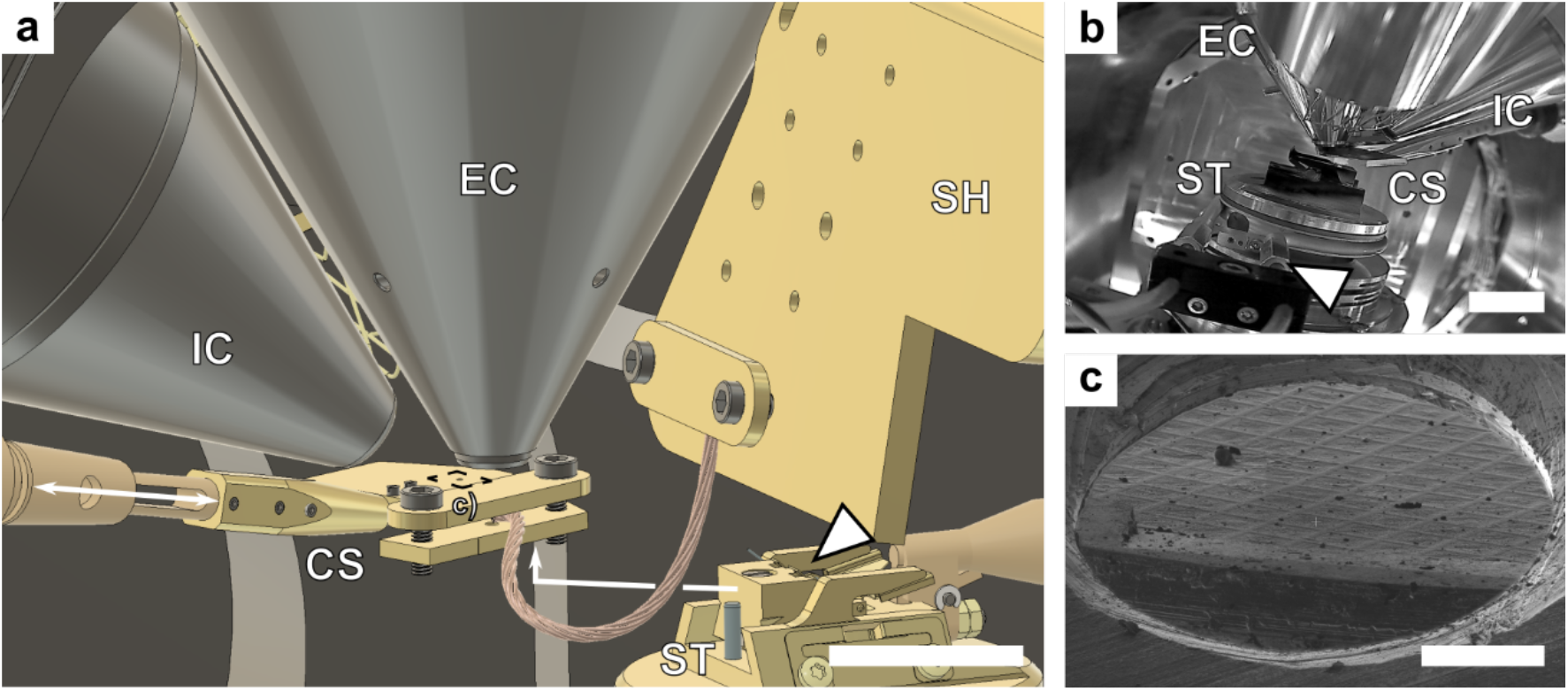
Cryo-shield and cryo-shutter. **a**) Schematic of the cryo-FIB/SEM chamber. The installed cryo-shield (SH) is installed next to the electron column (EC) and the ion column (IC). The total surface area of the cryo-shield (SH) is approximately 880 cm^2^. Assuming that the strength of the pump is ~ 15 l cm^−2^ s^−1^ [Bello2017], the estimated total pumping speed is ~ 13,000 l s^−1^. To reduce the partial pressure in the vicinity of the sample even further, a cryo-shutter (CS) can be inserted during milling (white arrow). The shuttle with the autoloader grids (white triangle) is moved via the cryo stage (ST) right below the cryo-shutter. Two different cryo-shutter types were tested (Supplementary Figure 1). Cryo-shutter Type II covers most of the grid reducing the contamination rate even further compared to the cryo-shutter Type I. Scale bar, 25 mm. **b**) Camera image from the back of the chamber. The cryo-shutter covers the autoloader grids almost completely. The white triangle indicates the position of the stage heater (not installed in this image). Scale bar, 25 mm. **c**) Ion image of the cryo-shutter Type II, showing the EM grid through the shutter hole. The diameter of the shutter hole is approximately 1 mm to ensure a complete protection of the grid. For electron imaging, the shutter is removed automatically. With these improvements, the sample can be prepared and stored in the milling position until imaging without amorphous ice contamination. Scale bar, 300 μm.

Due to the increased pumping speed, which is the strength of the cryo-pump, the partial pressure decreases from 2-3 x 10^−7^ mbar to approximately 4 x 10^−9^ mbar when the cryo-shield is cooled with liquid nitrogen. After approximately 30 min, the system reaches the lowest partial pressure of water (Figure 3b). If the cryo-shutter is inserted (Figure 4a,b), the partial pressure is reduced even further in the direct vicinity of the sample and under these conditions, the measured contamination rate is (5.6 ± 1.0) nm h^−1^, (Supplementary Table 1). Although this reduction is already remarkable, we refined the design of the cryo-shutter (Supplementary Figure 1). We minimized the diameter of the ion beam opening (Figure 4c), and completely removed the electron beam opening (Supplementary Figure 1). With the new design, the contamination rate was not measurable anymore (0 ± 2) nm h^−1^ (Supplementary Table 1, Supplementary Figure 2). It should be mentioned, that the initial contamination rate is different for each microscope. Typically, the contamination rate for the used cryo-FIB/SEM is specified to be smaller than 50 nm h^−1^. In combination, the improved vacuum conditions allow for milling over a longer time course and pave the way for supervised automated milling strategies.

### Automated lamella production pipeline

The lamella production workflow can be separated in four major steps: i) definition of region(s)-of-interest (ROI), ii) site preparation, iii) rough milling and iv) polishing (Supplementary Figure 3). In this study, we used the software application MAPS (Thermo Fisher Scientific) to visually identify ROIs directly in the cryo-FIB/SEM (Supplementary Figure 3a). If it is not possible to visually identify the ROIs or rare biological events, such as for instance cell-cell fusions, are investigated, the correlation of cryo-fluorescent light microscopy and cryo-FIB/SEM should be used. The information of the ROIs is directly transferred to the software application AutoTEM Cryo (Supplementary Figure 3b), where we set the parameters relevant for milling such as ion beam currents, dimensions of the milling pattern, and desired lamellae dimensions. The eucentric height and the milling angle are determined automatically by the program and rough milling, polishing, as well as the milling of micro expansion joints (relief cuts) to release the stress of the support layer [Wolff2019] are then executed in a fully automated manner (Supplementary Figure 3b, 4). This process resulted in up to 27 lamellae per session (16 h) with a final thickness of 100 - 200 nm (Supplementary Figure 3d).

### Efficiency estimation and quality assessment

To estimate the efficiency of our lamellae preparation pipeline, that is the production of thin and intact lamellae, we separated the handling steps into two parts: i) the pre-processing and ii) the post-processing. The pre-processing includes the assembly of the autoloader grid, its loading to the cryo-FIB shuttle and the transfer of the shuttle from the glove box to the cryo-FIB. The post-processing step comprises the transport from the cryo-FIB/SEM to the glove box and the transfer of the autoloader grid from the cryo-FIB shuttle to the autoloader cassette.

In recently published preparation protocols, the success rate of obtaining suitable specimens for proficient users was estimated to be 80% for the pre-processing and 90% for the post-processing step, which results in a total efficiency of 72% [Medeiros2018]. In reality, the efficiency is much lower because some lamellae are usable for cryo-ET analysis.

Our improvements of the pre- and post-processing steps resulted in a considerably higher efficiency exceeding 96% throughout the entire process (pre- and post-processing). During pre-processing no grid was lost and only 7 out of the 198 lamellae cracked during the post-processing. In addition, all intact lamellae were almost contamination free (Figure 5a). Applying the improved hardware, on average, only 9% of the lamella areas were covered with ice particles (Figure 5b) compared to 20% using the standard set up (Figure 5b, Figure 1a, Table 1). Importantly, in line with the fact that contamination inside the cryo-FIB/SEM was below the detection limit, we could not identify any contamination layer in the final tomograms (Figure 5c) and approximately 90% of all intact lamellae were useable for cryo-ET investigations (Table 1). Taken together, the high efficiency and tremendously reduced contamination rate reflect the impact of the newly implemented hardware components.

**Figure 5:**
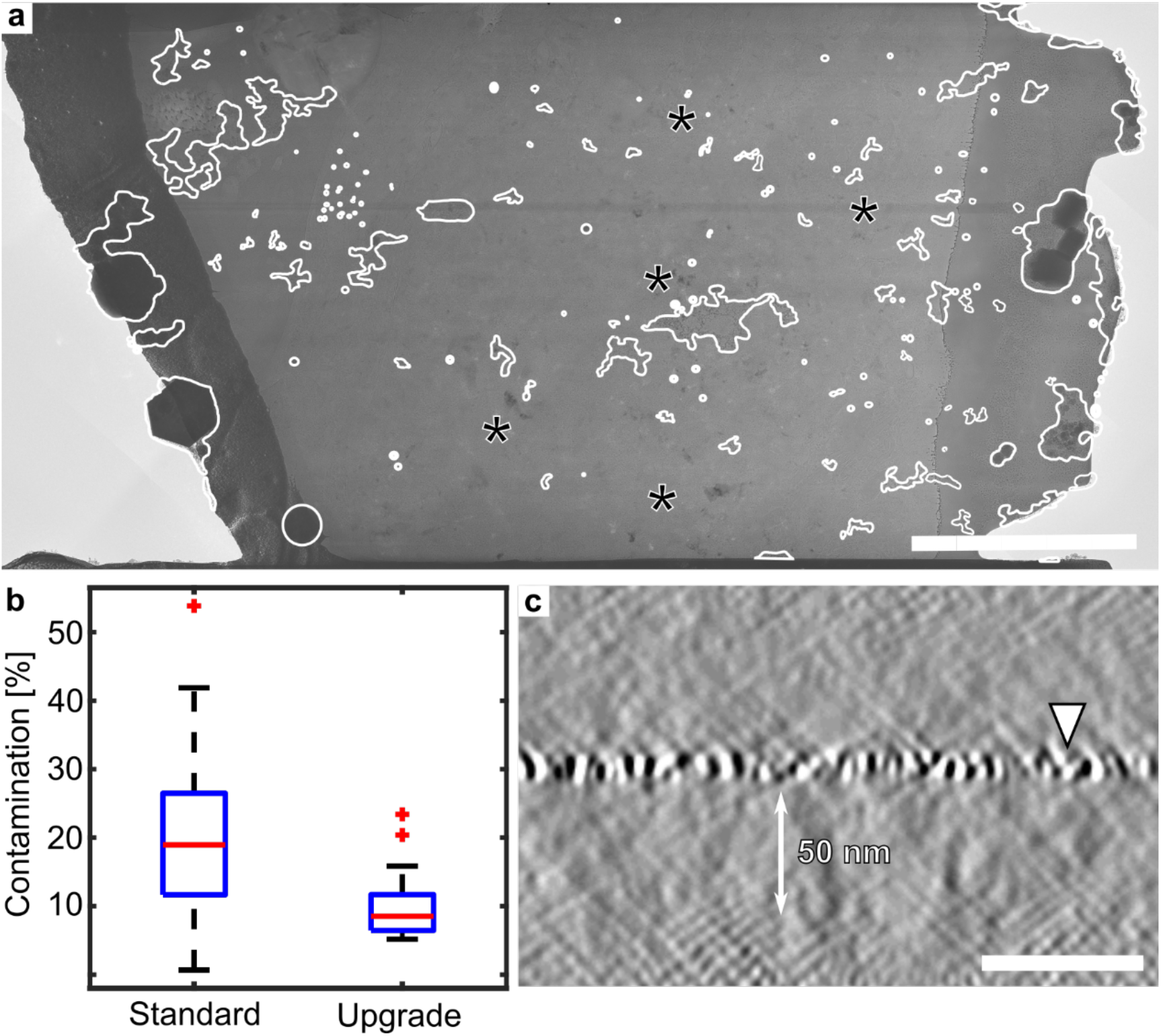
Quality assessment of the pre- and post-processing step. **a**) Overview of a lamella prepared with all hardware upgrades in use. In this case, only 8% of the lamella surface were covered with frost contamination. Most of the ice crystals are small enough that data acquisition is still possible. In some areas, non-vitrified regions can be identified (asterisks). Scale bar: 5 μm. **b**) Boxplot diagram of the frost contamination analysis. We analyzed 3000 μm^2^ of lamellae produced before and after our hardware upgrades revealing that the average frost contamination was on average reduced from 20% to 9% after installation of the new components. **c**) Side view of a lamella, prepared with all hardware upgrades in use. As indicated (white triangle), the sputter layer is directly on top of the lamella, indicating that no contamination occurred during milling. Scale bar: 50 nm.

**Table 1:** Impact of the hardware modifications on the frost contamination. To determine the impact of the hardware upgrades, the frost contamination without and with hardware upgrades was determined (Figure 5b). For each condition, three milling session were performed. An overview of each lamella was taken and the frost contamination was determined manually. Finally, the contaminated and total area were summed and the ratio was calculated.

| Experiment (sample) | Hardware upgrades | Number of ROIs | Successfully milled lamellae | Max. contamination thickness [nm] | Average area covered by ice crystals |
| --- | --- | --- | --- | --- | --- |
| Myofibrils | no | 18 | 12 | 50 | 20 % |
| Myofibrils | no | 11 | 6 | 100 |  |
| Cardiomyocytes | no | 11 | 8 | n.a. |  |
| **Success rate** |  |  | **65 %** |  |  |
| Myofibrils | yes | 11 | 11 | 0 | 9 % |
| Myofibrils | yes | 16 | 14 | 0 |  |
| Cardiomyocytes | yes | 7 | 4 | 0 |  |
| **Success rate** |  |  | **85%** |  |  |

To assess the quality of the automated lamellae preparation, we applied the automated lamella production pipeline as described above. We investigated different types of specimens and quantified the success rates for the rough milling and polishing. We considered a milling task as successfully completed, if the following criteria were fulfilled: i) the lamella was intact, i.e. not broken, ii) the desired thickness was reached, iii) tracking of the lamella site was successful, i.e. the right position of the sample was milled. We determined an average success rate of 96% and 88% for the rough milling and polishing, respectively (Table 2). With the estimated efficiency of 96% for the pre- and post-processing and the calculated efficiency of 84% for the milling process, the efficiency of the total workflow can be estimated to be around 80%. Although the success rate was much higher in the case of manual milling (above 95%), the total throughput is much higher, and could be further increased with overnight runs. Moreover, the operator time is significantly reduced.

**Table 2:** Milling statistics for different types of sample material using the automated lamella production pipeline.

| Sample | Number of ROI (lamellae sites) | Number of successfully roughly milled lamellae | Number of successfully polished lamellae |
| --- | --- | --- | --- |
| Myofibrils | 56 | 53 | 43 |
| Cardiomyocytes | 18 | 18 | 12 |
| A9 cells | 7 | 6 | 6 |
| *S. cerevisiae* | 94 | 89 | 85 |
| *E. coli* | 23 | 23 | 21 |
| Neurons | 3 | 3 | 2 |
| HeLa cells | 6 | 6 | 6 |
| Sum | 207 | 198 | 175 |
| **Success rate** |  | **96%** | **88% (total 84%)** |

To demonstrate that lamellae, prepared by automated cryo-FIB milling using AutoTEM cryo, are suitable for subtomogram averaging, a set of ribosomes were identified on several lamellae (Figure 6a,b). With only 15,000 particles, a structure of 14 Å (0.143 FSC criteria) was calculated (Figure 6c, e). Since the orientations and locations can be precisely determined, we were able to identify polysomes (Figure 6d).

**Figure 6:**
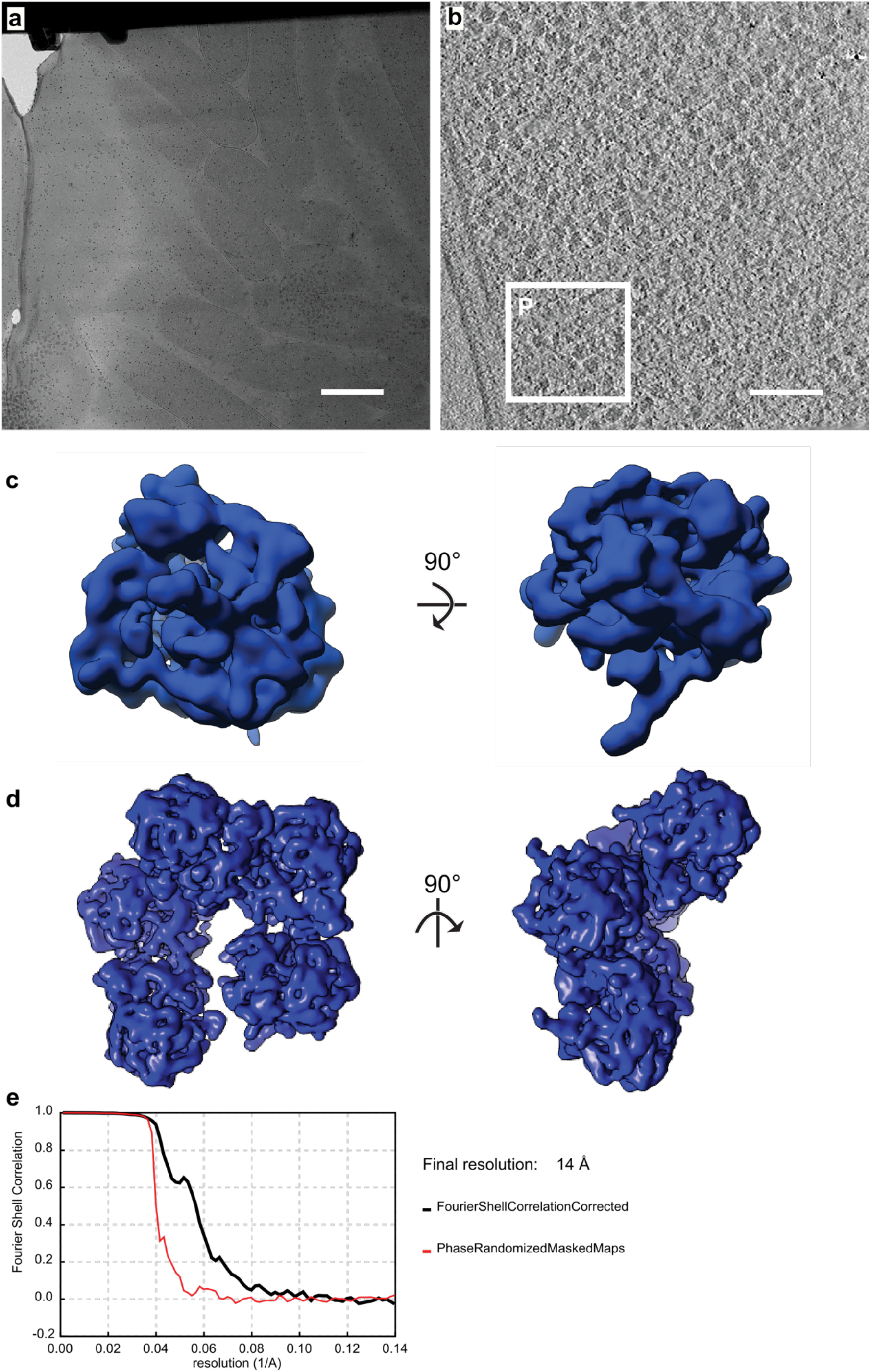
Suitability of automatically milled lamellae for subtomogram averaging. **a**) Representative overview of an automatically milled lamella (*E. coli*). The lamella was sputter coated to reduce charging artefacts. Scale bar, 2.5 μm. **b**) Macromolecular complexes such as ribosomes are readily detectable. Scale bar, 100 nm. **c**) Subtomograms can be averaged to high resolution (14 Å at 0.143 FSC criterion). **d**) Additionally, orientations and locations can be precisely determined, which enabled the detection of polysomes up to a pentamer (insert and boxed region P in (**a**)). **e**) FSC-curve of the final ribosome reconstruction.

Taken together, these results clearly demonstrate that the streamlined workflow not only increases the overall throughput of the lamellae production but also generates results that are suitable for in-situ high-resolution investigations.

## Discussion

Contamination of the samples and inconvenient sample handling are the dominant bottlenecks of current cryo-FIB workflows, which can decrease the quality of the final lamellae and limit the overall throughput per milling session. In the present study we overcame the aforementioned limitations by the introduction of new hardware components (Table 3) and show that the new hardware in combination with automatic cryo-FIB milling increases the overall throughput of high-quality lamella production.

**Table 3:**
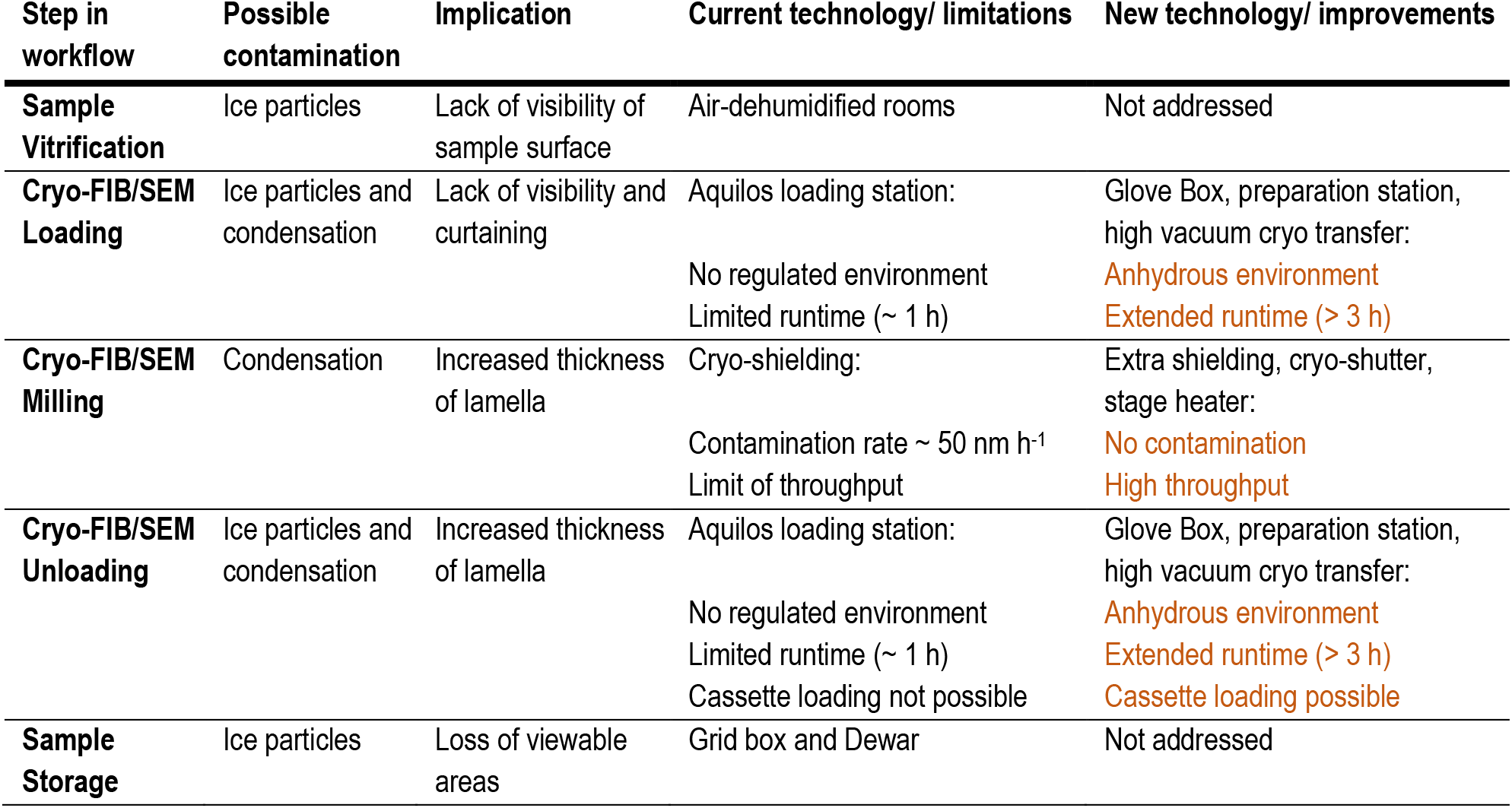
Overview of the improvements of the cryo-FIB/SEM workflow.

To reduce frost contamination throughout the entire preparation process, we introduced a glove box and a high vacuum cryo transfer system. Moreover, we reduced the amorphous ice contamination rates inside the vacuum system to a non-measurable level, mainly because of the installation of an additional cryo-shield and a cryo-shutter. Now, the entire handling and transfer steps as well as the cryo-FIB milling can be performed in an almost anhydrous environment. In order to streamline the handling of the grids carrying fragile lamellae, we introduced a new preparation station, which aims at avoiding all types of adverse environmental implications related to sample handling with maximal convenience for the user.

The contamination rate of 50 nm h^−1^ in the initial cryo-FIB/SEM setup limited the total number of prepared lamellae per session. Due to the improved and new hardware, we could prevent contamination in the milling position. Therefore, the lamella production is not restricted to a specific time frame anymore and, if a suitable automated filling system is in place, an 24/7 operation is even possible now.

We tested the new software application AutoTEM Cryo, which successively processes a queue of lamella sites in a supervised to automated manner, allowing for automatic batch milling of many lamellae. With the help of AutoTEM Cryo, the time spent on the microscope is now reduced to approximately 5 minutes per lamella site at the beginning of the milling process. Because of the automation of Aquilos cryo-FIB milling, we can now fully exploit the benefits of the hardware improvements and can indeed prepare up to 27 lamellae per session. All lamellae preparations in the present study were performed in sessions with an operation time of 8-16 hours.

In addition, the supervised automated milling process allows users to explore the influence of different milling parameters and strategies in a systematic manner. Due to the higher throughput, quality assessment routines can be developed and once these routines are established, the influence of different milling parameters can more readily be investigated, for example the influence of high ion currents (2-3 nA) on the quality of the cryo-lamella, which is not yet fully understood.

Without the contamination layer the tracking of the ROI position can be more easily determined by the AutoTEM Cryo software application. This became especially apparent in the case of myofibrils that due to their limited width proved to be a challenging sample for cryo-FIB milling. The improved conditions in the FIB-SEM tremendously increased the success rate of obtaining useable lamellae from ~ 65% to ~ 85% (Table 1).

Taken together, the combination of new hard- and software increases the overall throughput of lamellae production, improves the quality of the lamellae, and simplifies the process of cryo-FIB milling and sample transfer. In addition, the hardware components described in this work enable new types of experiments, which were previously difficult to realize.

## Data availability

The major hardware components developed in this study, such as the glove box, preparation station, and cryo shutter (Type II) can be purchased at Delmic B.V.

## Acknowledgements

We thank Sagar Khavnekar and Florian Beck for help with subtomogram averaging. This work was supported by funding from the Max Planck Society (to S.R. and J.P.). M.G. is supported by an EMBO Long-Term Fellowship.

## Material and Methods

### Glove box, preparation station and high vacuum cryo transfer

The glove box has been manufactured by Microscopy Solutions Py. Ltd and modified by us accordingly. Before sample preparation, the glove box was purged overnight with dry nitrogen (10 ppm H_2_O). Before the sample was brought into the glove box through the load lock, the liquid nitrogen reservoir and the preparation station were filled with liquid nitrogen. Then, EM grids were clipped to the autogrids, if not already done. Thereafter, the Autogrids were loaded into the cryo-FIB shuttle and transferred to the cryo-FIB/SEM (Aquilos, Thermo Fisher Scientific) via the high vacuum cryo transfer system. After the milling session was started, both the glove box as well as the preparation station were baked out at 40-50 °C with the help of the installed heaters. After milling, the cryo-FIB shuttle with the sample was transferred back to the glove box and the autoloader grids were unloaded, rotated by 90° and mounted to the autoloader cassette. Finally, the cassette was loaded into the NanoCab (Thermo Fisher Scientific) and transferred to the electron cryo microscope (Titan Krios, Thermo Fisher Scientific).

### Cryo-FIB/SEM

The developments described in this study were carried out on an Aquilos cryo-FIB/SEM (Thermo Fisher Scientific), equipped with a larger cryo-shield (manufactured by CryoVac GmbH & Co KG), a cryo-shutter (in-house production) and stage heater (in-house production). The instrument was equipped with a gas injection system (GIS) and a platinum sputter coater (Thermo Fisher Scientific). After insertion, the grids were coated with an organometallic platinum layer to protect the sample while milling. The sputter coating was utilized to avoid charging effects during imaging in the SEM and can either be done before and/or after GIS deposition. For scanning electron imaging, an acceleration voltage of 2-5 kV with an electron current of max. 28 pA was used. The ion optics were operated at 30 kV acceleration voltage with different ion currents (50 – 700 pA). All ion currents were specified by the user and should be tuned for every new sample material. The alignment of the ion optics is crucial and was performed at least once a week to avoid massive image shifts while changing the ion currents. For scanning electron imaging, the installed Everhart-Thornley detector was utilized.

### Automated milling via AutoTEM Cryo

Although the relevant milling parameter as ion beam currents, milling patterns and milling time might vary for different types of sample, a brief overview is given:

After the visual identification of ROIs via the software application MAPS (Thermo Fisher Scientific), the lamella sites were imported to the software application AutoTEM Cryo (Thermo Fisher Scientific). Thereafter, the automated milling routine was started. During the first step (preparation), the eucentric height and the milling angle were determined automatically. During the final action of the preparation step, the user was able to adjust the final milling position. In total, the preparation step takes approximately 2-3 min for each lamella site. Thereafter, relevant milling parameters were defined or loaded from a former template. The lamella width is mostly defined by the type of sample material and the condition of the specific ROI. Lamella widths of 8 – 10 μm were typically used. The desired lamella thickness was project dependent: in some cases, the project required a lamella thickness below 150 nm and in other cases, a thickness of 250 nm was still within the required limits. For the first estimation of milling times, a total sample thickness of 15 μm was assumed.

In the first milling step, the chunk milling, relief cuts were milled to release the stress from the support film of the grid, which might occur during plunge freezing. Here, a width of 1-3 μm and a height of 10-15 μm were typically used. The relief cuts were milled at relatively high ion beam currents (0.5–1 nA) for approximately 3-5 min.

Thereafter, most of the bulk material was removed during rough milling. This task is separated into four different milling steps. During the first milling step, most of the sample material was ablated with relatively high ion beam currents (0.5–1 nA) until a lamella thickness of approximately 2-3 μm was reached. In the following milling steps, the ion currents were consecutively reduced from 0.5 to 0.1 nA. After these milling steps, the lamellae thickness was already reduced to 300-600 nm. For the initial rough milling the front and rear pattern height were set to 5-7 μm. All later milling pattern were calculated accordingly. Since most of the sample material was removed during rough milling, most of the time was spent during this step (20-30min).

The polishing step was executed in three successive milling steps with an ion beam current of 30-50 pA. Since most of the sample material was already ablated, the last polishing process took only approximately 10 - 15 min.

### Cryo-electron tomography and tomogram reconstruction

All data sets were collected on a Titan Krios (Thermo Fisher), operated at an acceleration voltage of 300kV. The microscope was equipped with a field emission gun, a BioQuantum imaging filter (Gatan, Munich, Germany) with a set energy width of 20eV and a K3 direct electron detector (Gatan, Munich, Germany). The software package SerialEM was used for automated data acquisition [Mastronade2005]. For reconstruction of the tomograms, the software package IMOD was used [Kremer1996]. Imaging conditions are listed in Supplementary Table 2.

### Template Matching, Polysome Detection and Visualization

Tomograms were binned (IMOD bin 8x, 14.32 Å pix^−1^) and template matching was performed using PyTom [Hrabe2012]. First, a reference was constructed from approximately 300 manually picked ribosomes, which were aligned in PyTom using the fast-rotational matching (FRM) algorithm. The reference obtained from this was then truncated at the small subunit to allow later removal of false positives. For each tomogram, the 800 highest-scoring cross-correlation peaks were extracted, and the subtomograms aligned and classified using a maximum likelihood approach in Relion 3 [Zivanov2018] to remove false positives. This was judged by the absent small subunit in the false positive class averages. After initial processing at bin 8x, further classification and refinement of the 2x binned (3.58 Å pix^−1^) subtomogram averages was performed in Relion, including normalization and CTF estimation with CTFFIND 4.1.5 [Rohou2015]. The final list of particles (approximately 15000) yielded a ribosome structure at 14 Å resolution (0.143 FSC criterion), which was used in all animations. Polysome detection was performed analogously to published using an inhouse Matlab script [Brandt2010].

### Frost Contamination Measurements

To quantify the reduction of frost contamination, three lamellae preparations were performed with the standard configuration of the cryo-FIB/SEM and three preparations with the developed hardware upgrades (glove box, preparation station and high vacuum cryo transfer). After milling, overview images (montages) of each lamella were taken with the TEM. Each overview was analysed regarding the frost contamination by outlining each contamination spot manually (Figure 1a, Figure 5a). Only contaminations in direct contact with the lamella were taken into account. To avoid user bias, any contamination, regardless of its impact on the final data acquisition, was used for the analysis. Finally, the ratio of contaminated area and total lamella area was calculated (Figure 5b). The total lamellae area and the contaminated area were summed and the ratio was calculated for both conditions. For both conditions, a total area of approximate 3000 μm^2^ were analysed (Table 1). Whereas the experiments with the standard configuration were performed with a room humidity of approximately 30%, the experiments with the hardware upgrades were performed with a room humidity of 50-60%.

### Amorphous Ice Contamination Measurements

To quantify the impact of the hardware, which was installed inside the cryo-FIB/SEM (cryo-shield, cryo-shutter and stage heater), the growth of the amorphous ice contamination was measured. One day before the measurement, plain grids (Quantifoil 2/2) were transferred to the microscope. Directly before the measurement, the stage was moved into the milling position and thereafter, the cryo-FIB/SEM was cooled. One hour after cooling, the baseline images were taken. The next image was taken after at least two hours. For imaging, the sample stage was rotated by 110° and tilted by 30°. The following imaging conditions were used for all measurements: Acceleration voltage: 5 kV, working distance: 7mm, electron current: 25 pA, exposure time: 1 μs, pixel size: 1 nm, detector: T2 A+B, imaging mode: *optitilt*. For each measurement, two to three positions in different areas of the grid, including at least four holes each, were utilized. For every hole, the baseline thickness and the thickness with the gained contamination were determined (Supplementary Figure 2) and corrected for its residual tilting angle. Finally, all measurements of the specific condition were averaged.

---

**Supplementary Figure 1:**
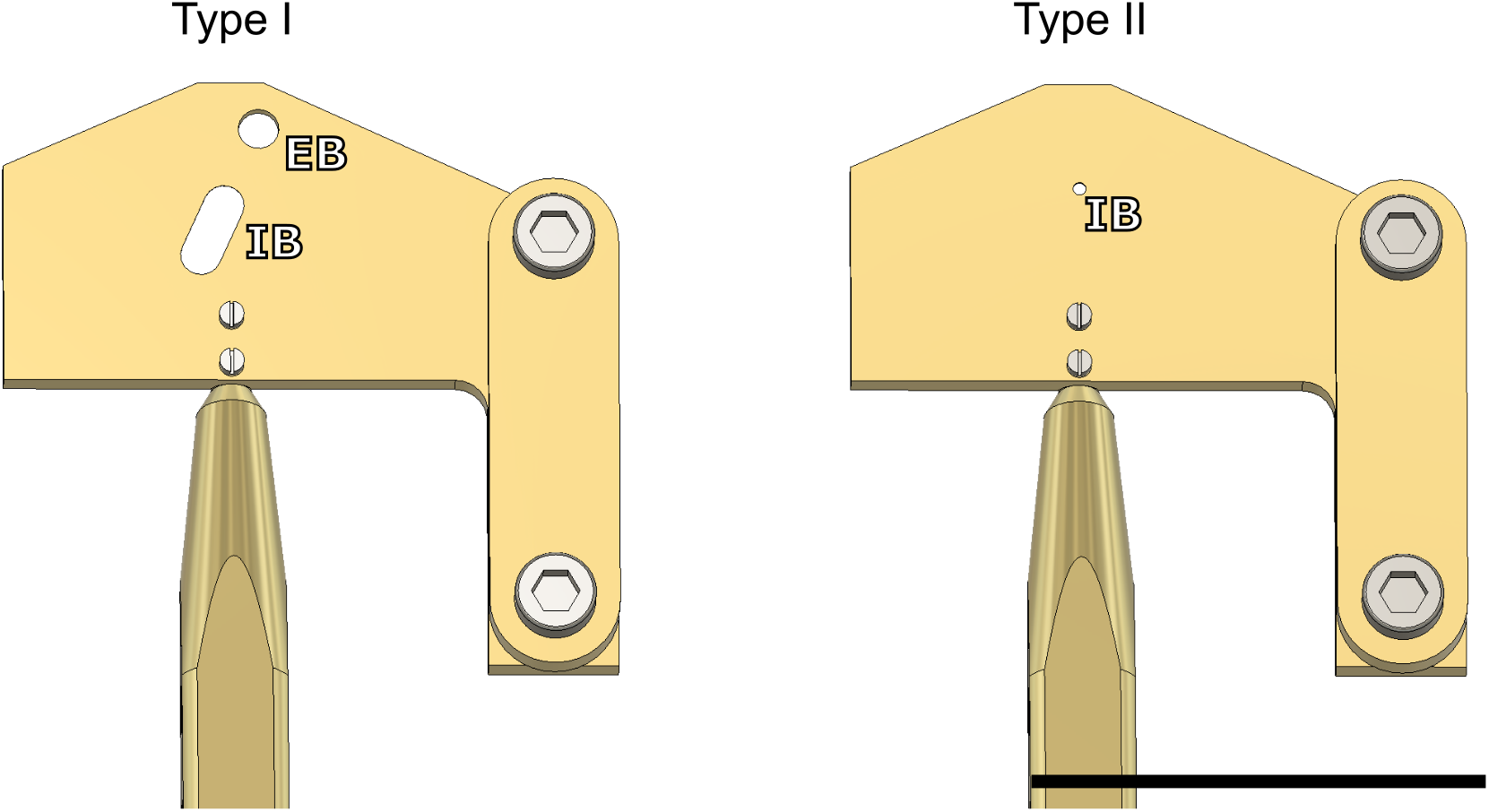
Cryo-shutter designs. For Type I, an opening for the ion beam (IB) and the electron beam (EB) were considered. To reduce the contamination even further, the opening for the ion beam was minimized as much as possible in the cryo-shutter Type II. Moreover, the opening for the electrons was removed completely. With these modifications we could reduce contamination to a non-measurable level. Scale bar, 28 mm.

**Supplementary Figure 2:**
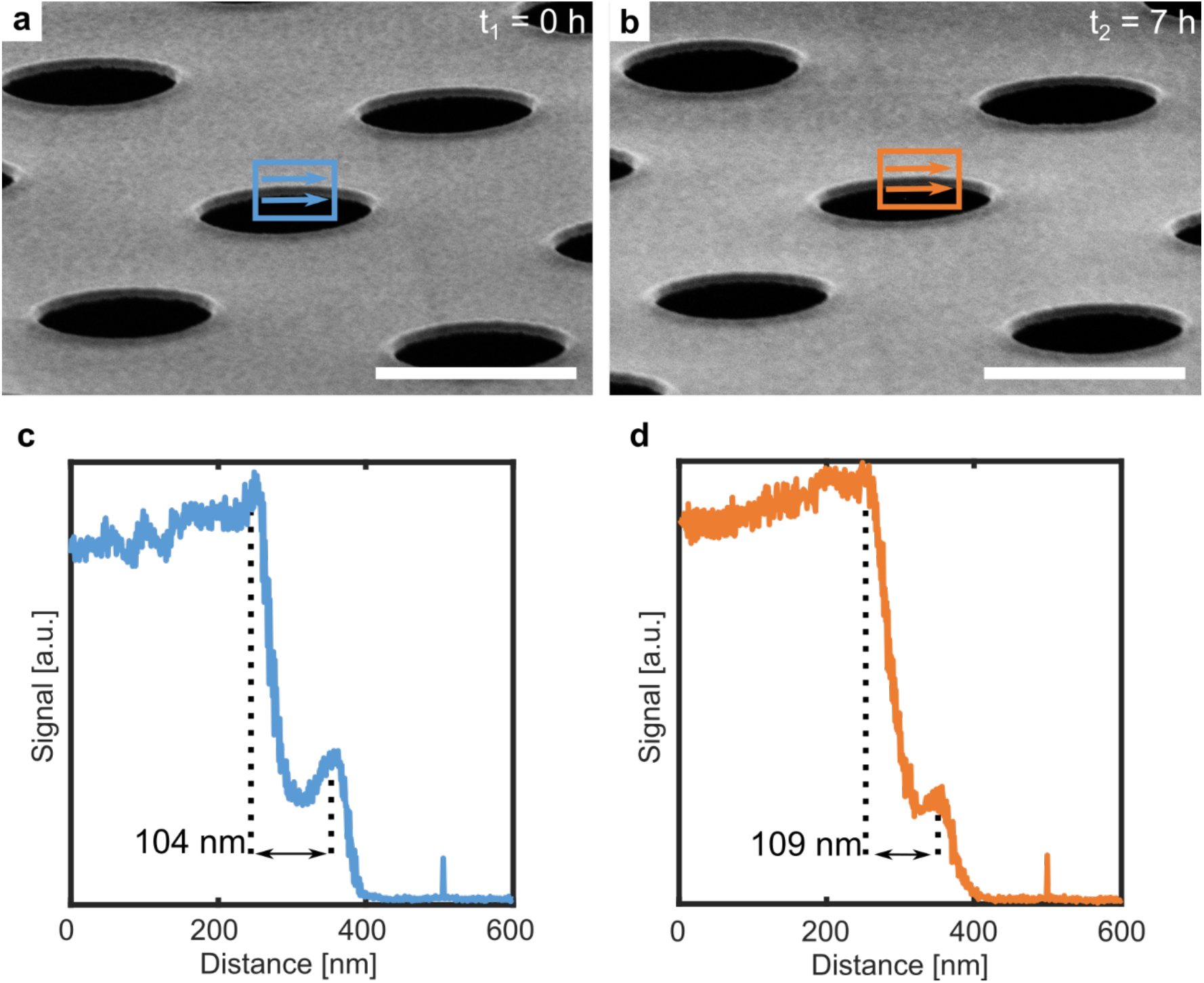
Measurement of the amorphous ice contamination growth rate in the milling position. **a**) Before imaging, the stage, cryo-shield and the cryo-shutter were cooled. After reaching the final temperature of approximately −185°C, the stage was heated to −165°C. Since there might be a temperature offset of 15°C between stage and shuttle, we assume that the sample has a temperature of approximately −150°C. Whereas the cryo-shield reached a temperature of −185°C, the cryo-shutter could only be cooled to −177°C. To determine the amorphous ice contamination rate in the milling position, a holey gold EM grid (UltraAuFoil holey gold grid) was mounted to the cryo-FIB/SEM shuttle and loaded into the instrument. For the baseline measurements (t_1_) the thickness of the holey gold film was determined (blue frame). To avoid ice ablation during focusing, beam tuning was done next to the region-of-interest. **b**) After 7 hours (t_2_), another series of thickness measurement was performed on the same holes. **c**) Baseline measurements: Line profiles were averaged horizontally (blue arrows in a)). The upper and the lower edge of the profile (dotted black line) were fitted with Gaussian function. If fitting was not possible, edge positions were determined manually. The centers of the edges were used to calculate the thickness. **d**) Thickness determination after 7 h: The line profile was averaged and the thickness was determined as described in c). Finally, both values were corrected regarding the tilting angle of the grid and the total increase in thickness was determined. Herewith, the contamination rate can be calculated. Scale bars, 2 μm.

**Supplementary Figure 3:**
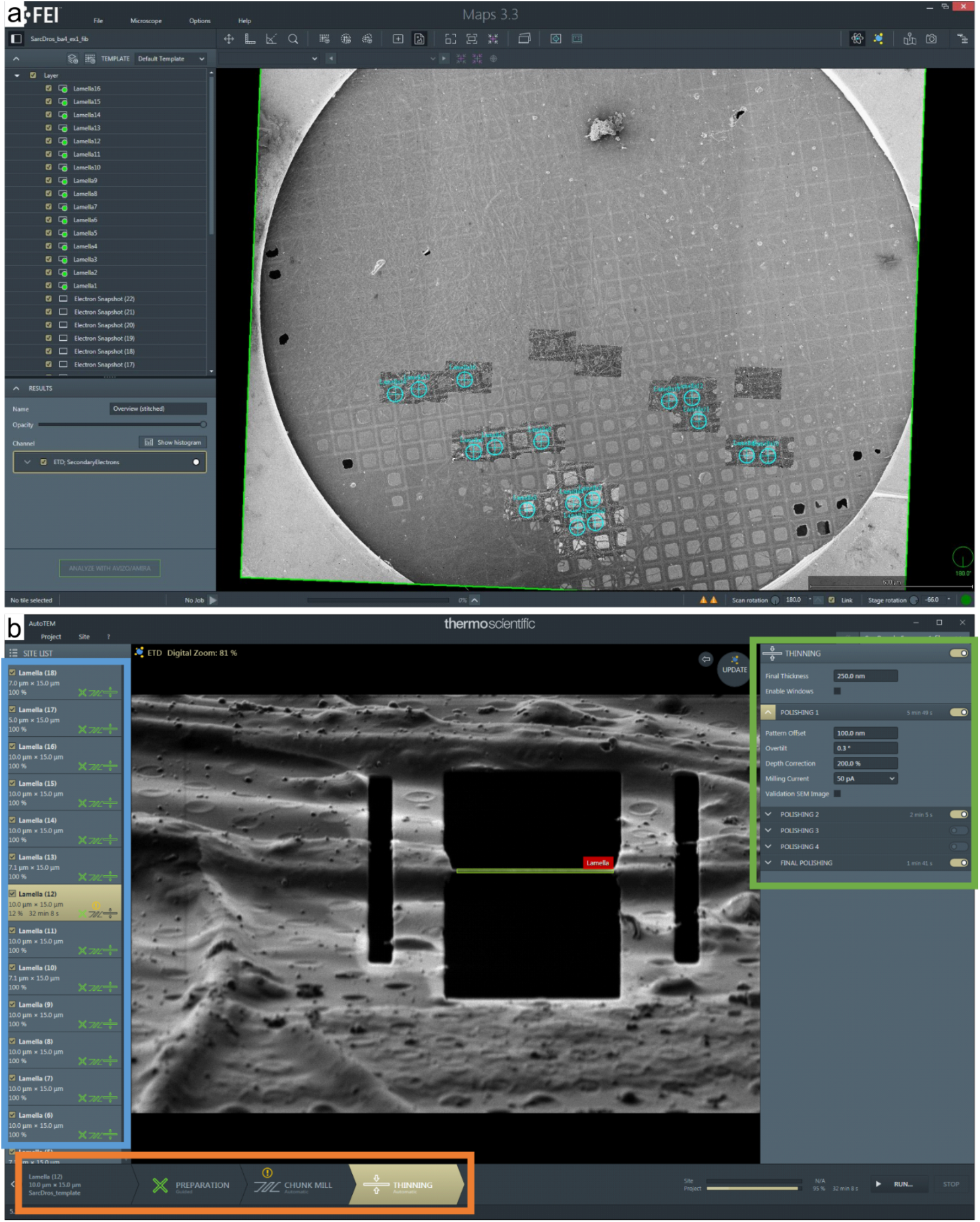
Software applications. **a**) Overview of the MAPS software application. Either ROIs are identified by light cryo microscopy and uploaded in the MAPS software or identified directly in the overview image as shown here. **b**) Overview of the AutoTEM Cryo software application. The software pipeline is separated into three different tasks (orange frame): i) Site preparation, ii) rough milling iii) thinning. Each step consists of several sub-tasks (green frame) which can be tuned to guarantee an optimal performance for each lamella site (blue frame).

**Supplementary Figure 4:**
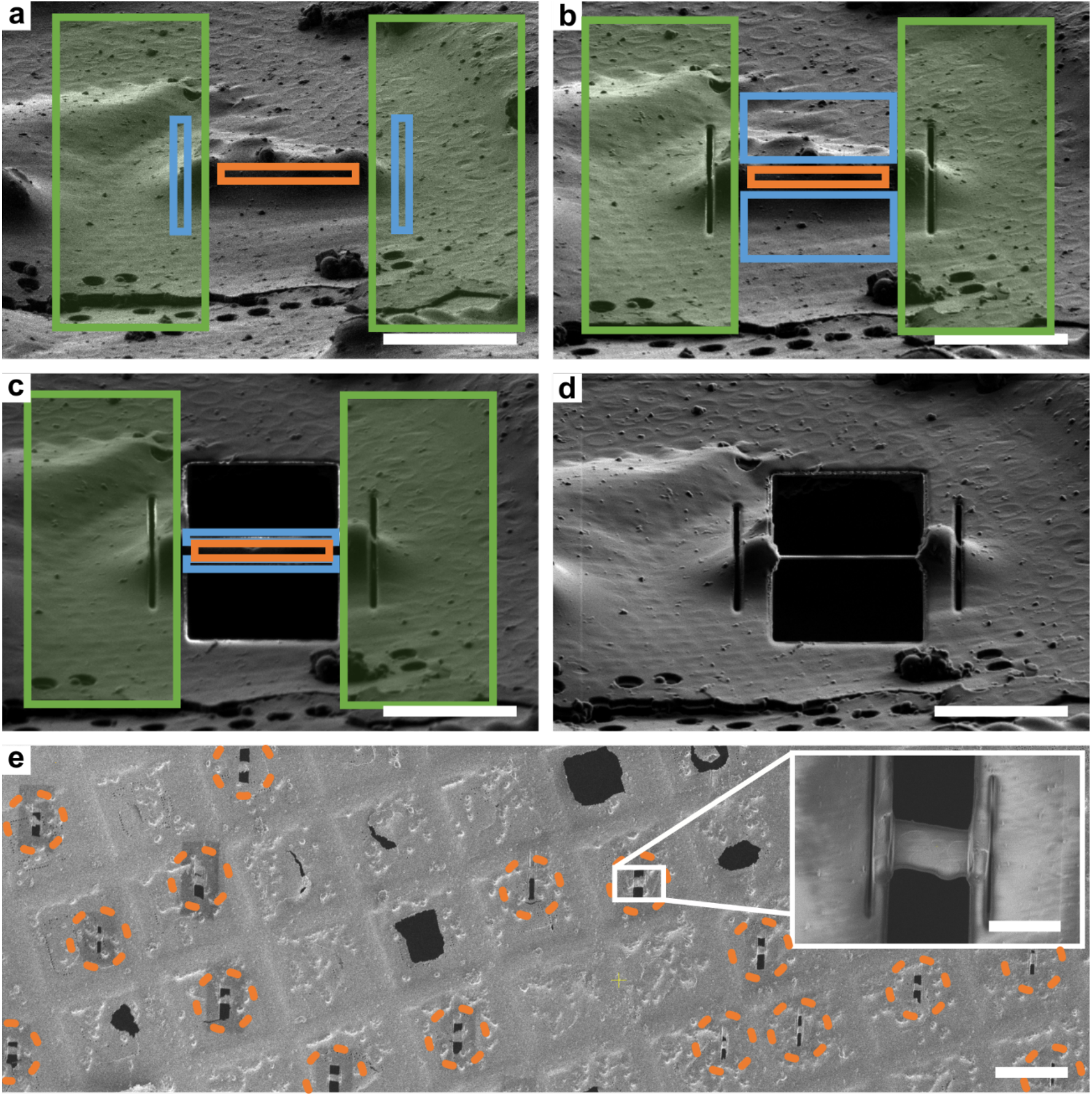
AutoTEM pipeline for automated lamellae preparation. **a**) After the lamella site is centered, micro-expansion joints (blue frame) are milled next to the lamella site to release possible stress of the support layer. Scale bar, 10 μm. **b**) Thereafter, rough milling (blue frame) is performed in four consecutive steps. First, the bulk material is ablated with relatively high currents (0.5 – 1 nA). In the subsequent steps, the lamella is thinned to a thickness between 250 – 500 nm with an ion current of 0.5 to 0.1 nA. This step is performed for all milling sites, before the final step is executed. Scale bar, 10 μm. **c**) In the last step, the lamella is polished (blue frame) with low ion beam currents (30 – 50 pA). Scale bar, 10 μm. In order to correct for possible stage drifts, specific regions of the ROI (green frames in a), b) and c)) are imaged frequently and compared by a newly developed, cross-correlation based algorithm. **d**) The final lamella thickness is optimally about 100 to 200 nm. Scale bar, 10 μm. **e**) Overview of the different lamella sites, prepared during the same session (dashed orange circles). Scale bar, 50 μm (Inset scale bar: 10 μm).

**Supplementary Table 1:** Influence of the different hardware components on the amorphous ice contamination rate inside the cryo-FIB/SEM. The number of holes, which were used for thickness determination (sample size), is given by the number *n*. For each condition (see also Supplementary Figure 1a), two to three positions, containing at least four holes, were selected. The uncertainty is estimated by the standard deviation of the different conditions.

| Condition | Stage temperature<br>[°C] | Shield temperature<br>[°C] | Status<br>shutter | Contamination rate<br>[nm h<sup>-1</sup>] (*n*) |
| --- | --- | --- | --- | --- |
| 1 | -188 | RT | Not inserted | 85 ± 7 (15) |
| 2 | -188 | -177 | Not inserted | 72 ± 4 (8) |
| 3 | -188 | -177 | Inserted | 12.8 ± 1.6 (15) |
| 4 | -165 | -177 | Inserted | 5.6 ± 1.0 (15) |
| 5 | -188 | -177 | Inserted | -0.6 ± 1.3 (15) |
| 6 | -165 | -177 | Inserted | 0 ± 2 (45) |

**Supplementary Table 2:** Imaging conditions for the presented data. The imaging conditions for the tomograms used for sub-volume averaging (shown in Figure 5) and the TEM image (shown in Figure 5) are listed.

| Sample | Defocus<br>[μm] | Tilting range<br>[°] | Increments<br>[°] | Pixel size<br>[Å] | Total<br>Dose<br>[e<sup>-</sup>/Å] |
| --- | --- | --- | --- | --- | --- |
| *E. coli* overview | 50 | 0 | 0 | 22.8 | 10 |
| *E. coli* tomograms | 1.5-3.5 | ± 60<br>dose symmetric | 3 | 1.79 | 120 |

## References

[Grange2017] Grange, M, Vasishtan, D., & Grünewald, K. Cellular electron cryo tomography and in situ sub-volume averaging reveal the context of microtubule-based processes. J. Struct. Biol. 197, 181–190 (2017).

[Bartesaghi2012] Bartesaghi A. et al. Secondary Structure Determination by Constrained Single-Particle Cryo-Electron Tomography. Structure 20, 2003–2013 (2012).

[Schur2016] Schur, F. K. M. et al. An atomic model of HIV-1 capsid-SP1 reveals structures regulating assembly and maturation. Science 353, 506–508 (2016).

[Vulovic2013] Vulovic, M. et al. Image formation modelling in cryo-electron microscopy. J. Struct. Biol. 183, 19–32 (2013).

[Al-Amoudi2004] Al-Alamoudi, A., Norlen, L.P.O., & Dubochet J. Cryo-electron microscopy of vitreous sections of native biological cells and tissues. J. Struct. Biol. 148, 131–135 (2004)

[Al-Amoudi2005] Al-Alamoudi, A., Studer, D. & Dubochet, J. Cutting artefacts and cutting process in vitreous sections for cryo-electron microscopy. J. Struct. Biol. 150, 109–121 (2005).

[Marko2007] Marko, M. et al. Focused-ion-beam thinning of frozen-hydrated biological specimens for cryo-electron microscopy. Nat. Methods 4, 215–217 (2007).

[Rigort2010] Rigort, A. et al. Micromachining tools and correlative approaches for cellular cryo-electron tomography. J. Struct. Biol. 172, 169–179 (2010).

[Hagen2015] Hagen C. et al. Structural basis of vesicle formation at the inner nuclear membrane. Cell 163 1692–1701 (2015).

[Mahamid2016] Mahamid J. et al Visualizing the molecular sociology at the HeLa cell nuclear periphery. Science 351, 969–972 (2016).

[Schaffer2017] Schaffer, M. et al. Optimized cryo-focused ion beam sample preparation aimed at in situ structural studies of membrane proteins. J. Struct. Biol. 197, 73–82 (2017).

[Medeiros2018] Medeiros, J.M. et al. Robust workflow and instrumentation for cryo-focused ion beam milling of samples for electron cryotomography. Ultramicroscopy 190, 1–11 (2018).

[Zach2020] Zach, T. et al. Fully automated, sequential focused ion beam milling for cryo-electron tomography. Elife 9, e52286 (2020).

[Buckley2020] Buckley, G et al. Automated cryo-lamella preparation for high-throughput in-situ structural biology. Journal of Structural Biology 210, 107488 (2020).

[Tacke016] Tacke, S. et al. A Versatile High-Vacuum Cryo-transfer System for Cryo-microscopy and Analytics. Biophysical Journal 110 4, 758–765 (2016).

[Echlin1992] Echling, P. Low-Temperature Microscopy and Analysis. Plenum Publishing Corporation, New York (1992)

[Umrath1983] Umrath, W.W. Berechnung von Gefriertrocknungszeiten für die elektronenmikroskopische Präparation. Mikroskopie 40, 9–34 (1983).

[Wolff2019] Wolff, G. Mind the gap: Micro-expansion joints drastically decrease the bending of FIB-milled cryo-lamellae. J. Struct. Biol. In Press (2019).

[Mastronade2005] Mastronarde D.N. Automated electron microscope tomography using robust prediction of specimen movements. J. Struct. Biol. 152, 36–51 (2005).

[Hrabe2012] Hrabe, T. et al. PyTom: a python-based toolbox for localization of macromolecules in cryo-electron tomograms and subtomogram analysis. J. Struct. Biol. 178, 177–188 (2012).

[Zivanov2018] Zivanov, J. et al. New tools for automated high-resolution cryo-EM structure determination in RELION-3. eLife 7, e42166 (2018).

[Rohou2015] Rohou, A. and N. Grigorieff, CTFFIND4: Fast and accurate defocus estimation from electron micrographs. J. Struct. Biol. 192, 216–221 (2015).

[Brandt2010] Brandt, F. et al. The Three-Dimensional Organization of Polyribosomes in Intact Human Cells. Mol. Cell 39, 560–569 (2010).

[Kremer1996] Kremer, J.R., Mastronarde, D.N. & McIntosh, J.R. Computer visualization of three-dimensional image data using IMOD. J. Struct. Biol. 116, 1–6 (1996).

[Bello2017] Bello, I. Vacuum and Ultravacuum: Physics and Technology. CRC Press (2017)

